# Focused X-ray Luminescence Computed Tomography using a Continuous Scanning Scheme

**DOI:** 10.1101/2021.02.04.429805

**Authors:** Michael C. Lun, Yile Fang, Changqing Li

**Author notes:** Corresponding Author: Changqing Li.

## Abstract

X-ray luminescence computed tomography (XLCT) imaging is a hybrid molecular imaging modality combining the merits of both conventional x-ray imaging (high spatial resolution) and optical imaging (high measurement sensitivity). The narrow x-ray beam based XLCT imaging has been shown to be promising. However due to the selective excitation scheme, the imaging speed is slow thus limiting its practical applications for *in vivo* imaging. In this work, we have introduced a continuous scanning scheme to acquire data for each angular projection in one motion, eliminating the previous stepping scheme and reducing the data acquisition time, which makes it feasible for multiple transverse scans for three-dimensional (3D) imaging. We have introduced a high accuracy vertical stage to our focused x-ray beam based XLCT imaging system to perform high-resolution and 3D XLCT imaging. We have also included a scintillator crystal coupled to a PMT to act as a single-pixel detector for boundary detection purposes to replace our previous flat panel x-ray detector. We have verified the feasibility of our proposed scanning scheme and imaging system by performing phantom experimental studies. A phantom was embedded with a set of cylindrical targets with 200 µm edge-to-edge distance and was scanned in our imaging system with the proposed method. To test the feasibility for 3D scanning, we took measurements from 4 transverse slices with a vertical step size of 1 mm. The results of the experiments verified the feasibility of our proposed method to perform 3D XLCT imaging using a narrow x-ray beam in a reasonable time.

## 1. Introduction

X-ray luminescence computed tomography (XLCT) was introduced in the past decade as a hybrid molecular imaging modality with great potentials for small-animal imaging by combining the high-spatial resolution of conventional x-ray imaging with the superb measurement sensitivity of optical imaging. Particularly, the narrow x-ray beam based XLCT has been shown to obtain very high spatial resolution, even at depths of several centimeters with good molecular sensitivity inside of turbid media [1, 2]. In principle, a focused or collimated beam of x-ray photons are utilized to penetrate deeply through the specimen with minimal scatter; any x-ray excitable contrast agents within the path of the x-ray beam will absorb the x-ray energy, causing a photophysical cascade of events leading to the emission of many optical photons which can pass through tissue and escape. Highly sensitive optical detectors such as an electron multiplying charge-coupled device (EMCCD) camera or photomultiplier tube (PMT) can then detect the emitted optical photons for image reconstruction. The first demonstration of XLCT imaging was reported by Pratx *et al*. using a selective excitation scanning scheme, much like first generation x-ray CT scanners, and demonstrated the potentials of this imaging method [3, 4]. Since then, due to several advantages of XLCT compared with other optical methods, several research groups, including our own, have pursued the improvement of XLCT from several aspects.

We have previously shown that by using a focused beam of x-rays as the excitation source for performing XLCT, orders of magnitude of increased sensitivity can be achieved due to higher flux and efficient use of x-ray photons compared with collimation. In addition, higher measurement sensitivity can also be obtained by using PMTs as the optical detector compared with EMCCD cameras [5]. While our results with this set-up are promising, the main drawback of this method is the relatively long measurement time due to the small beam size and selective excitation scanning scheme where the beam is moved or stepped along the object at predefined positions and the emitted photon intensity is acquired at each position before moving to the next position. To improve the scan time, in this work we have introduced a continuous scanning scheme where the x-ray beam will move across the object in a single continuous motion, and at predefined intervals, data will be acquired and saved by a high-speed oscilloscope such as the optical photon emission intensity from the object and the x-ray beam intensity by use of a single-pixel detector set-up used for boundary determination purposes. To accommodate three-dimensional (3D) data acquisition, we also introduced a motorized vertical lift stage in our set-up. To test the capabilities of our set-up, we have performed a phantom imaging experiment using the upgraded system.

The remainder of this paper is organized as follows: In Section 2, we discuss the upgraded design of our focused XLCT imaging system, the proposed continuous scanning scheme, and the experimental set-up. In Section 3, we show the results of our imaging experiment. Finally, in Section 4, we discuss our findings and conclude the paper.

## 2. Methods

### 2.1 XLCT experimental system set-up

Fig. 1 shows a computer-aided design (CAD) model of our proposed imaging system and Fig. 2 shows a photograph of the physical system in our laboratory. This imaging system is an upgraded version of the focused x-ray beam based XLCT imaging system previously described in [5]. In summary, an x-ray tube with a fixed polycapillary lens (X-Beam Powerflux [Mo anode], XOS) generates x-rays with a maximum energy of 50 kVp and tube current of 1.0 mA and are focused to an approximate focal spot size of 100 µm (focal distance: 44.5 mm). The imaged object is placed on a stage that within the focal spot of the x-ray beam and is fixed on top of a rotation stage (RT-3, Newmark Systems Inc.) mounted to a motorized vertical lift stage (VS-50, Newmark Systems Inc.) and linear stage (NLE-100, Newmark Systems Inc.) for rotating and translating the object at various depths. Compared with our previous imaging system in [3], the addition of a motorized vertical lift will accommodate feasible 3D scans since changing the imaging depth can now be automated instead of manually performed. The passed x-ray beam’s intensity is monitored by utilizing a single scintillator crystal mounted directly opposite of the beam and acts as a single-pixel detector. Emitted optical photons from this scintillator are collected by an optical fiber cable (labeled as Fiber 2 in Figs. 1 and 2) and then delivered to a PMT (H8259-01, Hamamatsu) connected to a high-speed oscilloscope (MDO3104, Tektronix). Monitoring of the beam intensity is used to determine the phantom boundary during measurements and replaces the previously used flat-panel x-ray detector, decreasing the size and price of the imaging system. During the XLCT scans, emitted optical photons from our imaged object that propagate to the surface are collected using a single optical fiber cable (labeled Fiber 1) and delivered to a fan-cooled PMT (H7422-50). The signal from the PMT is then amplified using a broadband amplifier (SR455A, Stanford Research Systems) with a gain of 125 and then filtered with a low-pass filter (BLP-10.7+, *f*_c_ = 11 MHz, Mini-Circuits) to reduce high-frequency noise before finally being collected by a separate channel of the high-speed oscilloscope. The entire imaging system up to the PMTs are placed inside of a light-tight and radiation shielding cabinet and controlled with a lab computer.

**Figure 1.**
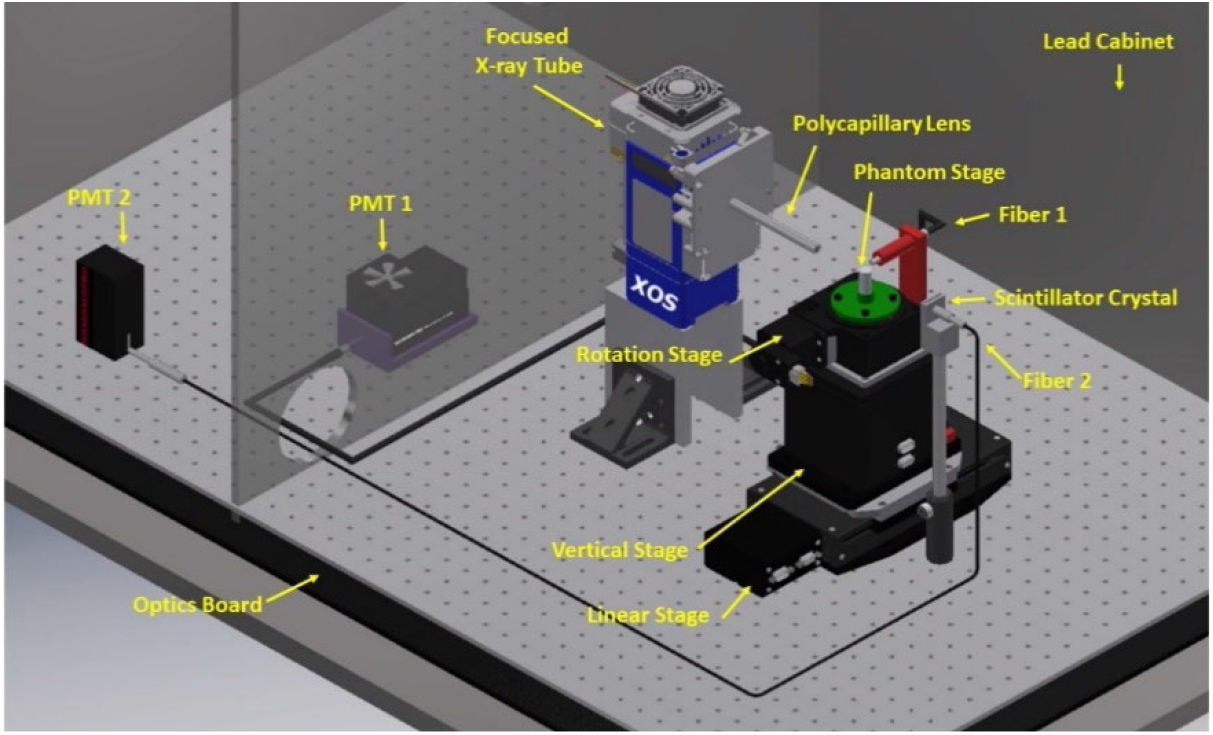
Computer-Aided Design (CAD) model of the 3D focused x-ray beam based XLCT imaging system.

**Figure 2.**
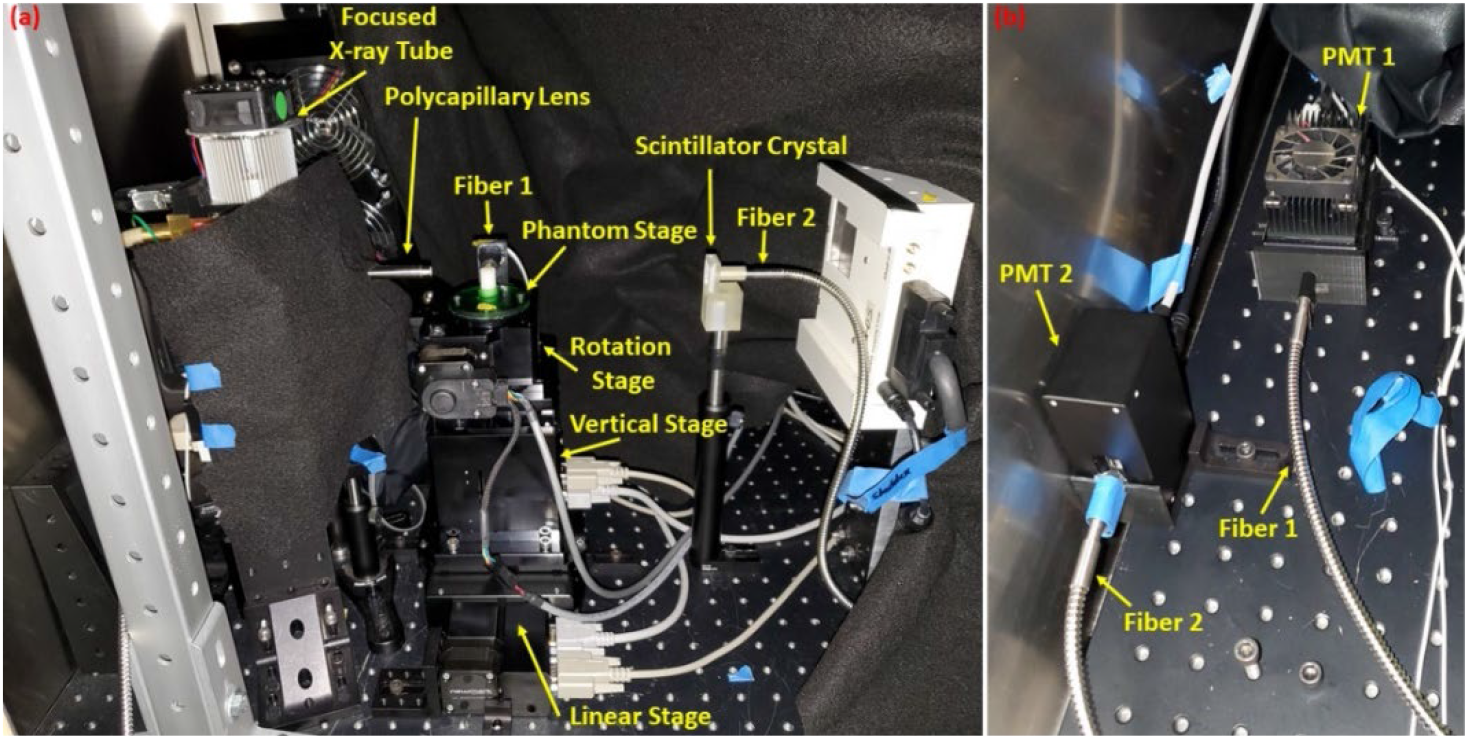
Photograph of the imaging system described in Fig. 1.

### 2.2 Scanning scheme

We developed a macro program written using Microsoft Visual Studios IDE in C++ to control and automate the XLCT scan. Previously, a conventional “step and acquire” scanning scheme was utilized during scanning for each angular projection. In this work, we implemented a continuous scanning scheme to significantly reduce the total scan time. Here, for each angular projection, a single movement of the linear stage is used to traverse the object. For example, for a 12 mm diameter, the linear stage will move continuously for 12 mm in one motion. Then by using information from the high-resolution encoder of the motorized linear stage, which is constantly monitored by our program, we can acquire and save data from two channels of the oscilloscope (corresponding to the emitted optical photons from the phantom and the scintillator) at defined intervals or spacings during the single stage movement. After each angular projection or linear scan, the next acquisition was acquired in the reverse direction (right to left, then left to right, etc.) to save time from having to reset the linear stage position. Also, for each consecutive slice or transverse section scanned, the rotation stage took measurements in the reverse direction than the previous to save time from resetting the stage. For each XLCT scan, we can input the total scanning distance (determined by the object size), interval spacing for data acquisition for each linear scan, linear translation speed which depends on parameters such as the interval spacing (finer spacing requires slower translation to save the data), the number of angular projections for each transverse section (in degrees), number of slices or transverse sections, and spacing between slices. To test the capabilities of our imaging system, we performed a phantom experiment as described next.

### 2.3 Phantom experimental set-up

A cylindrical phantom composed of 1% intralipid, 2% agar, and water with a diameter of 12 mm and height of 20 mm was created to be scanned in our system. Two glass capillary tube targets (outer diameter: 0.4 mm; inner diameter: 0.2 mm, Drummond Scientific) were filled with 10 mg/mL of Gd_2_O_2_S:Eu^3+^ (UKL63/UF-R1, Phosphor Techn. Ltd.) x-ray excitable contrast agent in a similar background solution as the phantom and embedded side-by-side in the phantom (edge-to-edge (EtE) distance of 0.2 mm). The phantom was then placed into the imaging system and scanned. During the XLCT scan, the x-ray tube was operated at a setting of 30 kVp and 0.5 mA and measurements were acquired from 6 angular projections (30°/projection). For each projection, the linear stage moved in one continuous motion for 12 mm and data was acquired from both channels of the oscilloscope every 0.15 mm for a total of 80 times each projection. The oscilloscope acquired a total of 10 ms of data from both PMTs for each acquisition. After all projections for a slice was finished, the vertical lift was then set to change the scanning depth by 1 mm and measurements were repeated. In total, we took measurements from 4 transverse slices at 4 different scan depths (defined as distance from the phantom top surface) of 5, 6, 7, and 8 mm, respectively. Following the XLCT scan, the phantom was then placed inside of our microCT scanner to perform a conventional CT scan. For the microCT scan, we acquired projection images using 180 projections of 2^°^step sizes. MicroCT images were reconstructed in MATLAB using a filtered back-projection algorithm with a Shepp-Logan filter. For XLCT imaging, image reconstruction is similar to fluorescence molecular tomography (FMT) [6]. Images were reconstructed using an optical photon propagation model (radiative transport equation) inside turbid media which also included information such as the x-ray beam’s size and location as spatial priors. We used the *L*^1^ regularized majorization-minimization algorithm developed originally for FMT but adapted to solve the XLCT inverse problem to reconstruct our XLCT images. Details of the algorithm are described in [7-10].

## 3. Results

Figure 3 below shows slices from the microCT reconstruction corresponding to the scanning depths for the XLCT scan. Using the microCT images, the center of the phantom was determined, then the distance from the phantom center to each target center was used as the ground-truth locations.

**Figure 3.**
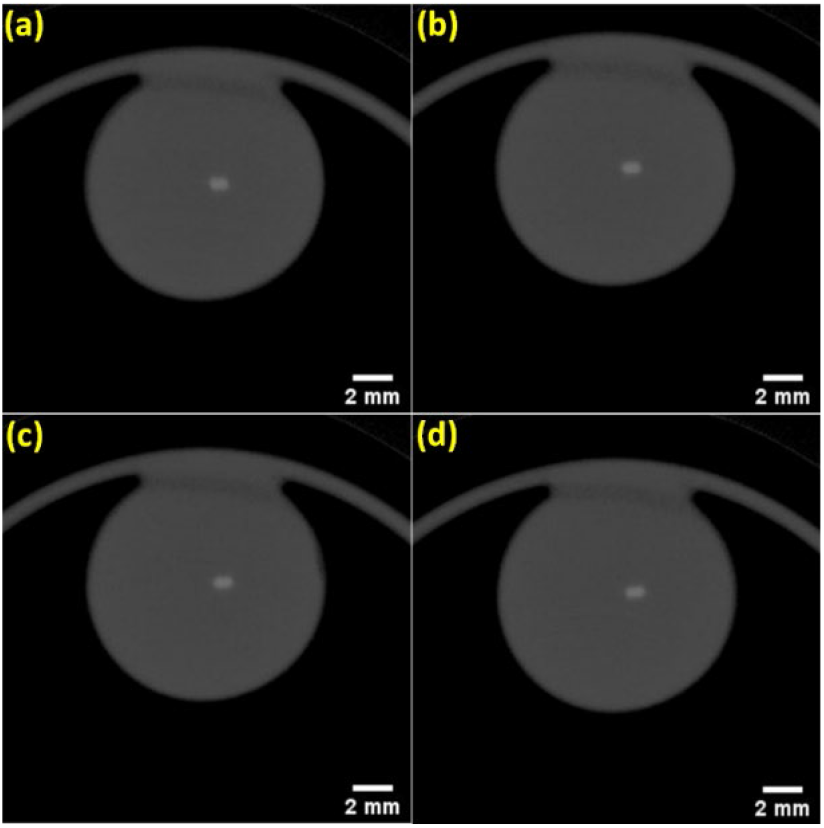
microCT reconstructed images of phantom used in experiment. Slices corresponding to depths of (a) 5 mm, (b) 6 mm, (c) 7 mm, and (d) 8 mm.

For the XLCT reconstructions, the scanned sections were interpolated onto a fine 2D grid with pixel size of 25 µm^2^. The system matrix was interpolated onto the grid from a system matrix calculated on a finite element mesh. Reconstructed XLCT images for the different scanned sections are shown below in Fig. 4 and the zoomed-target regions are shown in Fig. 5 where the green circles represent the ground truth obtained from the microCT reconstructions. For all four scanned sections, we can see that the two targets have been successfully resolved and are reconstructed at the correct locations. To quantitatively evaluate the reconstructions, the DICE similarity coefficients (using full width 10% maximum) is calculated for each reconstruction from which we achieved a DICE of 85.0, 77.3, 80.6, and 83.9 % for scan depths of 5, 6, 7, and 8 mm, respectively. Based on these metrics, we can see we have a high similarity between the reconstructed targets and the ground truth. In terms of the scan times to acquire the XLCT data, we were able to take measurements for each transverse section (with 6 angular projections) in 82 secs with a total scan time for all four sections totaling 338 secs (5 min and 38s).

**Figure 4.**
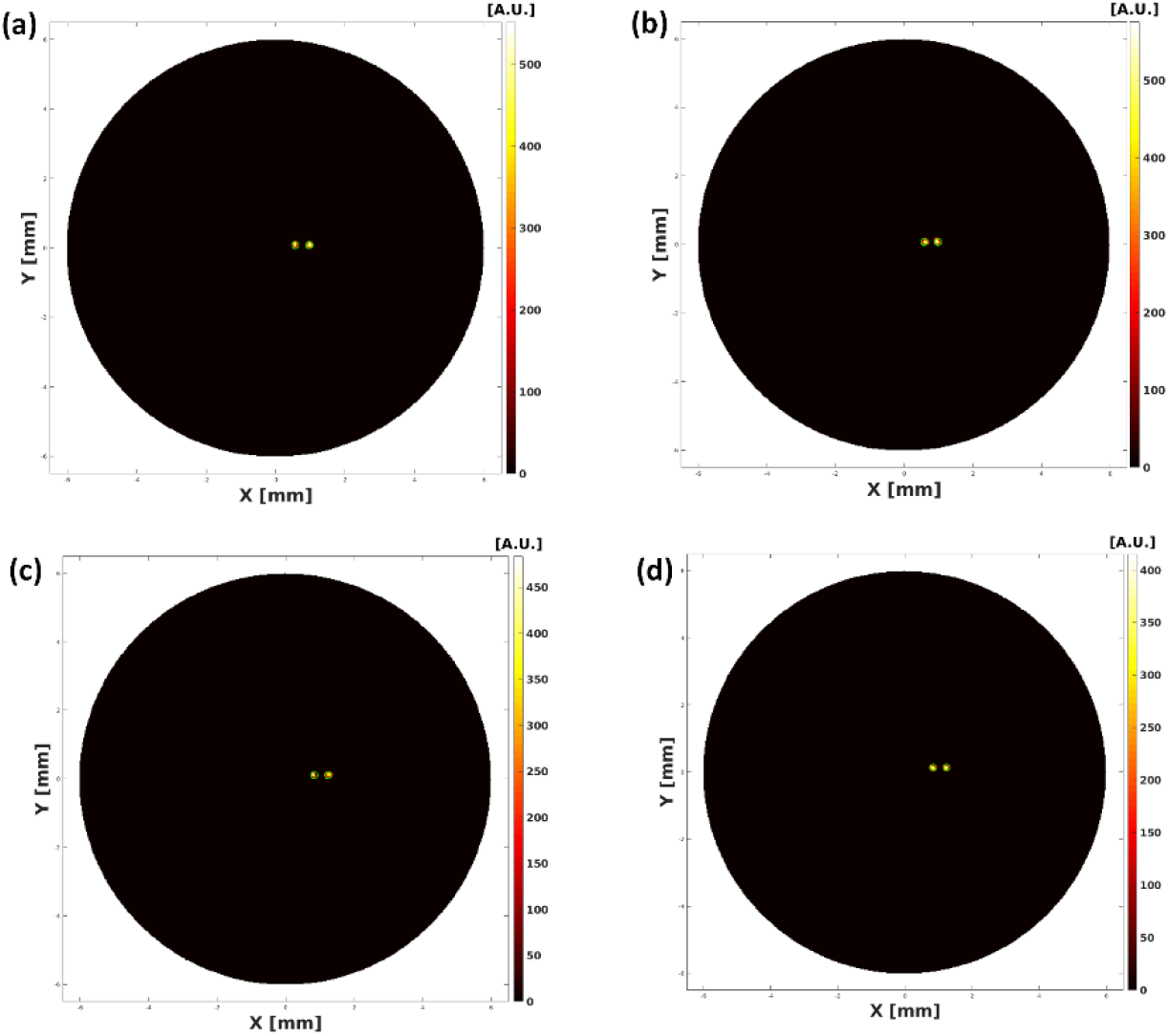
Zoomed-in reconstructed XLCT images for different scanning depths acquired during 3DXLCT scan. Scanning depths of (a) 5 mm, (b) 6 mm, (c) 7 mm, and (d) 8 mm. Green circles represent the ground-truth locations.

**Figure 5.**
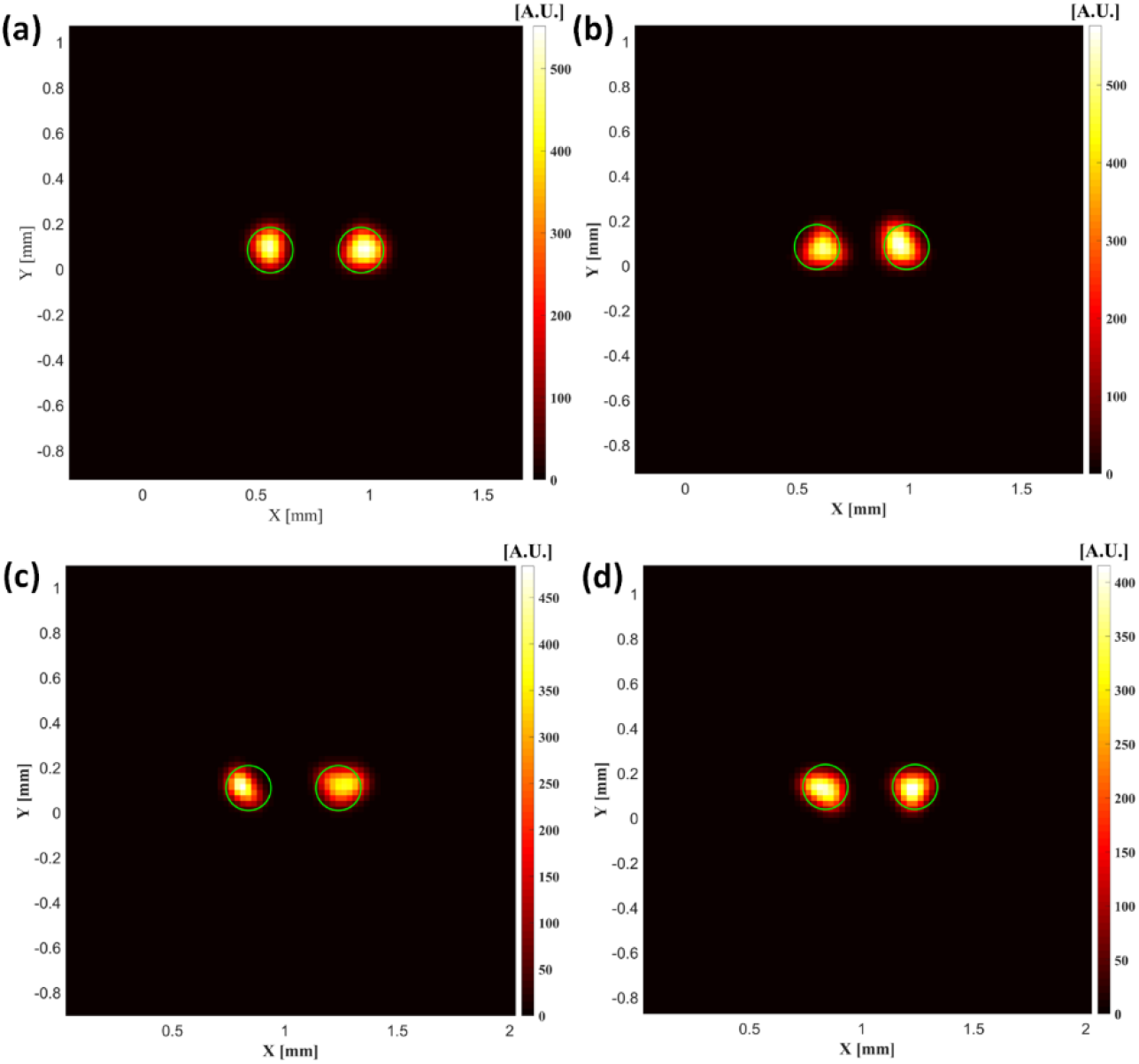
Zoomed-in reconstructed XLCT images from Fig. 4. Scanning depths of (a) 5 mm, (b) 6 mm, (c) 7 mm, and (d) 8 mm. Green circles represent the ground-truth locations.

## 4. Discussion and Conclusions

XLCT has emerged as a promising and attractive tool for small-animal molecular imaging. However, limitations due to the slow scanning speed hindered its applications for dynamic, *in* vivo, and whole-body imaging. In this work, we have proposed and investigated a continuous scanning strategy for narrow x-ray beam based XLCT imaging to improve upon the long imaging time. Compared with our previous studies, here we were able to perform a 6-projection transverse scan in 82 secs including all stage movements compared with 292.8 secs as in [11] (also including stage movements), an approximately 3.5 times reduction in scan time. Moreover, we were able to maintain the high-spatial resolution capabilities of this method and were able to successfully separate both targets with an EtE distance of 0.2 mm for all four cases while maintaining a good DICE similarity coefficient for all reconstructions as seen in Figs. 4 and 5. We have only performed XLCT scans from four transverse sections to demonstrate the capabilities of our current set-up. In future studies, we plan to investigate methods to further reduce the scan time. Currently a limitation to how fast we can perform our scans is the time necessary to save the data from the oscilloscope. If the movement speed of the stage is set too high, there is not enough time to save the data from the oscilloscope at the correct position as the stage is constantly moving. Currently it takes approximately 100-200 ms for data from both channels of the oscilloscope to save at each interval position. We are currently planning to use a dual-channel photon counter to replace the oscilloscope in our current set-up which will be a subject of future investigations.

## Acknowledgements

This work was funded by the NIH National Institute of Biomedical Imaging and Bioengineering (NIBIB) [R01EB026646].

